# Multiple Types of Context-Specific Brain Causal Regulatory Networks and their Applications to Autism Spectrum Disorder

**DOI:** 10.1101/2025.01.17.633619

**Authors:** Li Wang, Eric Yang, Gustavo Stolovitzky, Eric Schadt, Jun Zhu

## Abstract

One of the primary goals of systems biology is to understand how different cell types are regulated at various developmental stages and how dysregulation in these cells contributes to disease. With advances in high-throughput omics profiling technologies, vast amounts of both bulk tissue and single-cell data have been generated to study cellular regulation. To investigate the types of gene-gene regulatory interactions that can be uncovered from bulk versus single-cell profiling data, we constructed multiple context-specific brain gene regulatory networks using brain bulk and single-cell RNA-seq datasets and applied two distinct computational methods: an integrative Bayesian Network (BN) and a deep learning model, Geneformer. A total of 39 networks were generated, comprising 13 brain region-specific Bayesian networks from bulk RNA-seq data (bulk_BNs), 13 cell-type and developmental stage-specific Bayesian networks, and 13 corresponding Geneformer-based networks from scRNA-seq data (sc_BNs and sc_GFs, respectively).

We compared these networks in terms of their topological features, biological pathway regulation, and known regulatory relationships among disease-related genes. Our analysis revealed distinct network properties driven by data source type (bulk vs. single-cell RNA-seq) and network inference method (integrative Bayesian Network vs. deep learning Geneformer). Neurodegenerative disease-related pathways were co-regulated in bulk_BNs, while sc_BNs and sc_GFs captured pathways associated with neuroactive ligand-receptor interactions. To demonstrate the utility of these networks, we leveraged them to investigate potential disease mechanisms of Autism Spectrum Disorder (ASD). We found that ASD risk genes were enriched among hub genes common across networks but were more frequently hubs in context-specific networks, underscoring the significance of inhibitory interneurons during early developmental stages in ASD. Additionally, we showed that these networks could effectively recapitulate context-specific ASD gene signatures.

In summary, this study highlights the complementary value of integrating multiple data types through different computational approaches to advance our understanding of disease-related molecular mechanisms, such as those involved in ASD. Our findings provide a framework for using context-specific gene regulatory networks to identify disease-causal genes and key regulators of signaling pathways, potentially guiding future therapeutic strategies.

## Introduction

The brain is the most complex human organ, containing hundreds of cell types organized spatially into distinct compartments, each with specialized functions. Understanding how different cell types are regulated across developmental stages and how dysregulation in these cells contributes to disease is fundamental in biology. Recent technological advancements in RNA sequencing, particularly single-cell RNA sequencing (scRNA-seq) and spatial RNA sequencing, have enabled detailed profiling of the brain, covering different regions at both bulk tissue and single-cell resolutions^1-3^. Various methods, such as co-expression and causal network analyses, have been applied to bulk RNA-seq data to elucidate gene regulatory mechanisms underlying disease^4,5^. Diverse techniques have also been applied to scRNA-seq data to infer gene regulatory networks for individual cell types, aiming to understand cell type-specific regulations that govern diverse brain functions^6^. However, most network approaches used for scRNA-seq data reveal associations rather than causal relationships. In this study, we aimed to examine different causal network inference methods and explore the unique molecular insights that each type of network can provide.

The integrative Bayesian network (BN) approach has been applied to infer causal regulatory relationships that drive complex disease mechanisms^7,8^. While this methodology is well-established for bulk RNA-seq data, its utility for single-cell data remains underexplored. In contrast, deep learning models like Geneformer^9^ show promise in analyzing scRNA-seq data through in-silico gene perturbation studies. Comparing these methodologies in constructing context-specific causal regulatory networks can yield valuable insights into the strengths and limitations of each approach. To systematically assess networks inferred by these methods, we assembled a set of RNA-seq data profiling the human brain at both bulk tissue and single-cell levels. We constructed an ensemble of causal regulatory networks, comprising 13 tissue-specific Bayesian networks from bulk RNA-seq data, 13 cell-type and developmental stage-specific Bayesian networks from scRNA-seq data, and 13 cell-type-specific causal networks based on the Geneformer model. We then compared the structural and functional properties across these three sets of networks.

Autism Spectrum Disorder (ASD) is a complex neurological and neurodevelopmental disorder with a strong genetic component^10^. However, the primary causes, cell types involved, and underlying molecular networks contributing to disease onset and progression remain largely unknown. Over the past two decades, genomic research has identified hundreds of genes associated with elevated ASD risk. Translating these findings into a detailed understanding of ASD’s molecular mechanisms requires a deeper exploration of dynamic gene regulatory networks within the brain’s complex cellular environment. To demonstrate the utility of these networks in identifying disease-causal genes, we leveraged these context-specific networks to investigate how known ASD risk genes impact biological pathways across various cellular and developmental contexts. We illustrated how these network types can provide novel insights into ASD mechanisms, including the identification of key regulatory nodes and exploration of signaling pathways that may contribute to disease pathogenesis and serve as potential therapeutic targets. This study underscores the importance of integrating multiple data types and computational approaches to understand complex neurological diseases comprehensively.

## Results

### Constructing Tissue, Cell-Type and Developmental Stage-Specific Gene Regulatory Networks from Bulk and Single-Cell RNA-seq Profiles

Using an integrative Bayesian network approach^11^, we constructed 13 brain region-specific gene networks by integrating bulk gene expression and genetic profiles (eQTL) from GTEx consortium^1^, along with known protein complex, TF-target and pathway information (Methods, Figure 1A). The ages of GTEx subjects range from 20 to 79 years, with a median age of 50-59. After restricting to samples with paired bulk gene expression and genetic profiles, the sample size for each tissue ranged from 114 to 209, with a median of 170. Context-specific priors for protein complexes, TF-target and pathway-target pairs were incorporated into the network construction (see Methods for details).

**Figure 1.**
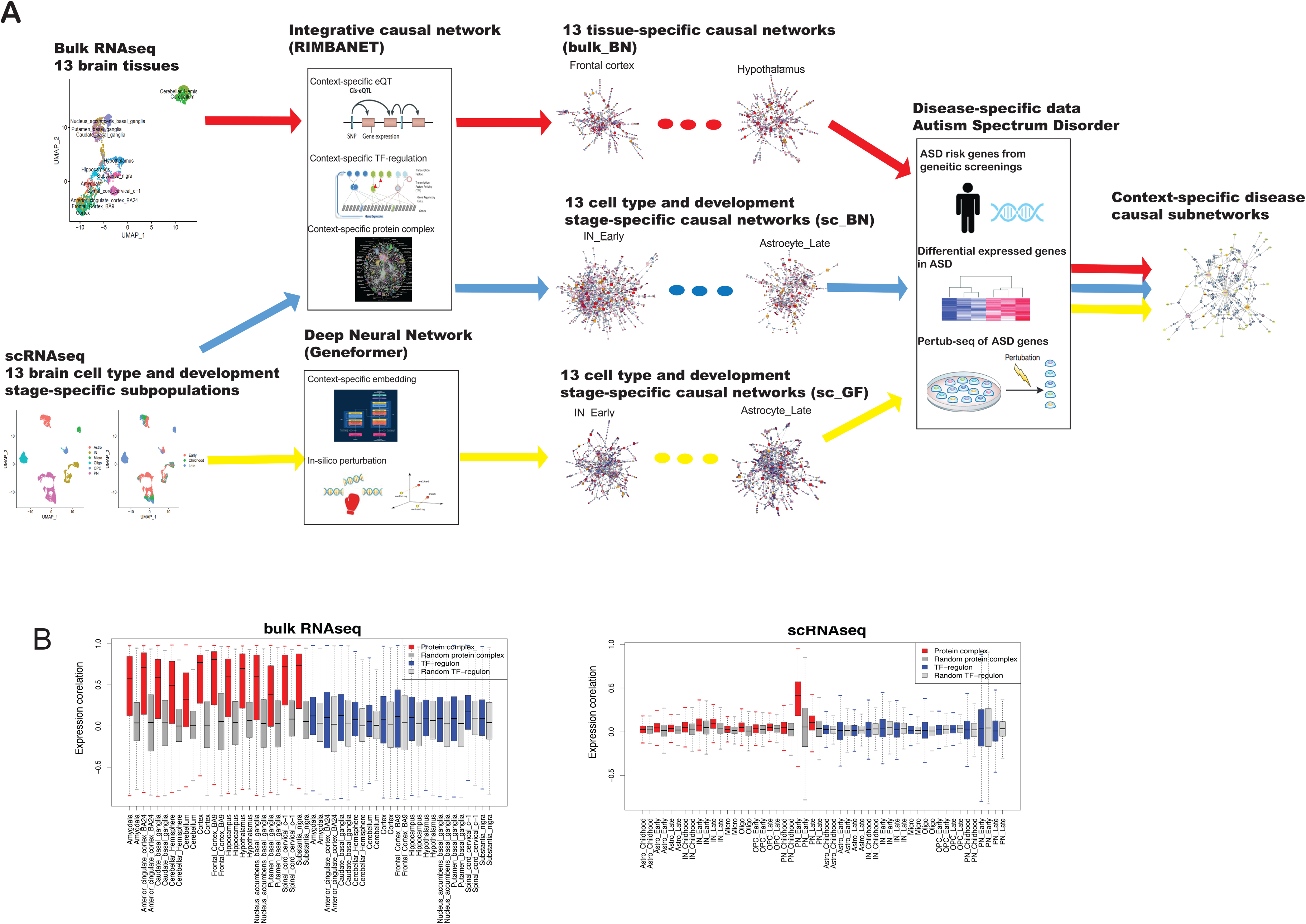
Overview of Context-Specific brain Causal Regulatory Networks. **(A)** The workflow for constructing context-specific causal regulatory networks based on bulk RNAseq and single-cell RNAseq (scRNAseq) profiles. Causal regulatory networks were constructed for 13 brain regions using bulk RNAseq and for 13 cell-type and development stage-specific subpopulations using scRNAseq. Two different computational approaches, the integrative Bayesian network (RIMBANET) and deep neural network (Geneformer), were employed to construct causal regulatory networks. Integrating those causal networks with disease specific data supports the identification of context-specific causal subnetworks underlying disease (e.g., ASD). **(B)** Comparation of expression correlations among members of known protein complexes and TF regulons for bulk RNAseq and scRNAseq profiles.

Using a similar integrative approach, we also constructed 13 cell-type and developmental stage-specific brain gene regulatory network from single nuclear RNA-seq(snRNA-seq) profiles (Methods, Figure 1A). The single-cell data encompass 152,250 cells (grouped into six major cell types) from 27 human prefrontal cortex samples^3^. When feasible, samples were divided into three developmental stages: early-stage (9 samples ranging from 22 gestation weeks to one year old with median of 86 days), childhood (8 samples of 1-8 years old, with a median of 2.5 years), and late-stage (10 samples from 10-40 years old, with a median of 16.5 years). To address the data sparsity associated with scRNA-seq profiles, meta-cells were generated by combining single cells with similar profiles (see Methods for details).

In addition to the Bayesian network approach for inferring gene-gene causal regulations, recent advancements in context-specific embedding within the deep learning field, such as the Geneformer algorithm^9^, show promising results for in-silico gene perturbation in a context-specific manner. Using the same brain snRNA-seq data described above, we fine-tuned the Geneformer model and conducted system-wise in-silico perturbations. The results were summarized into 13 cell-type and developmental stage-specific brain gene regulatory networks (Methods, Figure 1A).

In total, we constructed three sets of brain gene regulatory networks: 13 tissue-specific networks from bulk RNA-seq using the integrative Bayesian network approach (referred to as bulk_BN), 13 cell-type and developmental stage-specific networks from snRNA-seq using the integrative Bayesian network approach (referred to as sc_BN), and 13 cell-type and developmental stage-specific networks based on the Geneformer model (referred to as sc_GF).

### Network Topology Differences Reflecting Bulk vs Single-Cell RNA-seq Differences and Network Construction Method Difference

The number of nodes and edges in each of the 39 gene networks can be found in Table S1. The number of nodes ranges from 8,179 to 13,999 per network, and the number of edges ranges from 9,262 to 40,401. The node degree (the number of connections each node has) of these networks generally follows a log-normal distribution (Figure S1A). Node and edge counts, as well as the node degree distribution, were similar between sc_BN and bulk_BN networks, as the same gene selection and network construction method was applied to both. In contrast, the sc_GF networks exhibited higher numbers of nodes and edges, with node degree distributions that differed significantly from those of the other two network sets. These differences are mostly likely due to methodological differences between causal Bayesian network and the Geneformer approaches (See methods for details).

When comparing the average clustering coefficient of each network, bulk_BN networks had substantially higher value than sc_BN networks (Figure S1B). Given that the same construction method was used, the much higher clustering coefficient in bulk_BNs compared to sc_BNs likely reflects inherent differences between bulk and single-cell profiles, as well as distinct gene regulatory patterns they reveal. (For sc_GF networks, the clustering coefficient exhibited a wider range, with a median between that of bulk_BN and sc_BN. This larger range is likely due to the varying node and edge counts in sc_GF networks.)

Protein complex members in bulk profiles displayed a much higher expression correlation compared to randomly selected gene sets of the same size, but this difference was less pronounced in scRNA-seq profiles (Figure 1B). An exception was observed in the PN_Early group, where substantial gene expression changes in PN cells are expected during early development stage^3^. This disparity in correlation patterns contributed to noticeable differences in the number of structure priors for transcriptional co-regulations of protein complex components between the bulk_BNs and sc_BNs (Table S1), which in turn contributed to the higher clustering coefficients seen in bulk_BN networks.

On the other hand, genes within known TF-regulons generally did not show transcriptional co-regulation in either bulk (Figure 1A) or scRNA-seq data (Figure 1B). Since TF regulation is known to be context- and tissue-specific, this result suggests that the TF-target pairs in the curated database used here are not specifically brain-related. Notably, that number of structure priors for TF-target regulations incorporated in sc_BNs was similar to that in bulk_BNs (Table S1.csv).

### Enriched Connectivity Among Neuroactive Ligand and Receptor Genes in Gene Networks from scRNA-seq

We investigated nodes with high out-degree (hub nodes) in each network, as their perturbation is expected to produce widespread effects in the system. In particular, we focused on the commonalities and differences of hub nodes across the 39 networks. Approximately the top 500 high-degree nodes were selected from each network (using the largest degree cutoff that captures at least 500 nodes), and overlaps between hub nodes in each network pair were calculated using the Jaccard Coefficient. To minimize platform differences, we focused on 20,185 genes shared between bulk and scRNA-seq profiles. Each network set generally shared more hub genes within its own network type (Figure 2A). Interestingly, sc_BN networks appeared more similar to sc_GF networks than to bulk_BN, indicating the data source has a stronger influence than the specific algorithm used. Among the three sets, bulk_BN networks exhibited the highest similarity with each other, followed by sc_BN and then sc_GF, suggesting that the cell-type-specific regulatory patterns are more distinct than brain region-specific ones. The number of shared hub genes within each network set generally correlated with their profile similarity. For instance, the two cerebellar networks in bulk_BN shared fewer common hub genes with other tissue-specific bulk_BN networks, which is consistent with their RNAseq profiles shown in the UMAP plot (Figure S2A), Neuron-related networks (PN and IN) shared more hub nodes with each other than with non-neuron networks. Additionally, developmental stage variation was minimal between children and late-stage PN and IN networks, as reflected by the high overlap of hub nodes in sc_BN_PN_childhood vs. sc_BN_PN_late and sc_BN_IN_childhood vs. sc_BN_IN_late. In contrast, early-stage PN and IN networks exhibited more stage-specific regulatory patterns.

**Figure 2:**
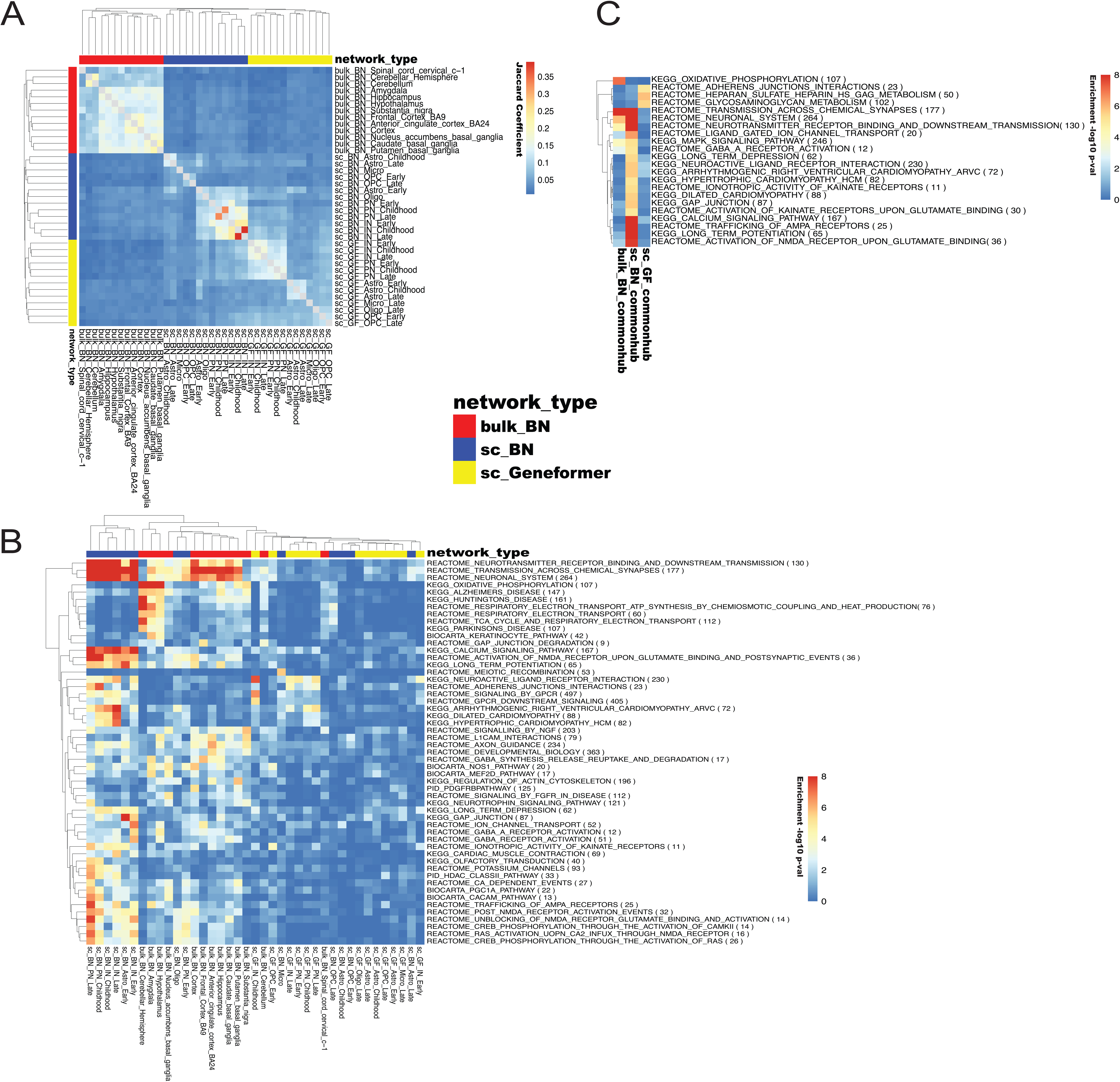
Comparative Analysis of Causal Regulatory Networks Across Bulk and Single-Cell RNA-seq Data. **(A)** Jaccard coefficient of hub gene overlap between bulk RNA-seq Bayesian networks (bulk_BN), single-cell RNA-seq Bayesian networks (sc_BN), and Geneformer-based networks (sc_GF). **(B)** Pathway enrichment analysis of hub genes for each context-specific network. The color gradient represents the enrichment score based on a hypergeometric test (-log10 p-value). For visualization purposes, the enrichment score has an upper limit of 8. **(C)** Pathway enrichment results for common hub genes shared within bulk_BN, sc_BN, and sc_GF networks, respectively. Color is the same as (B).

We then performed functional enrichment analysis on the hub genes in each network. As shown in Figure 2B, both bulk and single-cell networks were enriched with neural signaling pathways, such as REACTOME_TRANSMISSION_ACROSS_CHEMICAL_SYNAPSES and REACTOME_AXON_GUIDANCE. However, certain neuronal gene sets were more context specific. For example, KEGG_NEUROACTIVE_LIGAND_RECEPTOR_INTERACTION and REACTOME_ADHERENS_JUNCTIONS_INTERACTIONS were primarily enriched in hub nodes from scRNA-seq networks, but not in bulk_BN networks. Conversely, pathways related to KEGG_OXIDATIVE_PHOSPHORYLATION, KEGG_ALZHEIMERS_DISEASE, KEGG_HUNTINGTONS_DISEASE, and KEGG_PARKINSONS_DISEASE were enriched exclusively in hub nodes of certain tissue-specific bulk_BN networks. The lack of enrichment of neurodegenerative disease-related genes in sc_BN and sc_GF networks may be attributed to the younger donor age in our scRNA-seq dataset. Additionally, hub genes in sc_GF networks were enriched in fewer gene sets compared to those in sc_BN networks.

We further categorized the hub gene based on their frequency as hub nodes across the 39 networks. The distribution of hub node frequency followed a log-normal distribution (Figure S1C), with the majority of hub genes being context-specific and fewer hub genes shared across multiple networks. We defined common hub genes (1,233 genes) as hub nodes in at least five networks and classified the remaining 8,256 as context specific. Common hub genes were further clustered into three groups (Figure S1D): bulk_BN_common-hubs (573 genes), sc_BN_common-hubs (326 genes) and sc_GF_common-hubs (334 genes), corresponding largely to the network type they were most frequently associated with. Functional enrichment analysis revealed that bulk_BN_common-hub genes were uniquely enriched in oxidative phosphorylation while sc_BN_common-hub genes were uniquely enriched in several neural-specific pathways (Figure 2C).

Besides hub nodes, we also compared gene connectivity and functional enrichment across 39 networks. Gene connectivity in each network was represented by a vector of shortest path lengths (SPL) calculated for every gene pair (the vector length equals to the total number of gene pairs), and the network similarity was measured by the correlation coefficient of these SPL vectors. Network similarity based on SPL (Figure S1E) largely corroborated the observations from the hub gene analysis (Figure 2A). We further examined connectivity within each functional gene sets by calculating the average SPL for each gene set and comparing it to randomly selected gene set of the same size. This yielded a Z-score reflecting connectivity strength for each gene set. Figure 3A shows the top gene sets with differential Z-scores between bulk and single-cell networks. For example, NEUROACTIVE_LIGAND_RECEPTOR_INTERACTION (Figure 3B) and GPCR signaling pathways were more connected in sc_BN and sc_GF networks than in bulk_BN networks. In contrast, OXIDATIVE_PHOSPHORYLATION (Figure 3C) and inflammation-related pathways (e.g., TNF-related complex) were more connected in bulk_BN networks compared to sc_BN or sc_GF.

**Figure 3:**
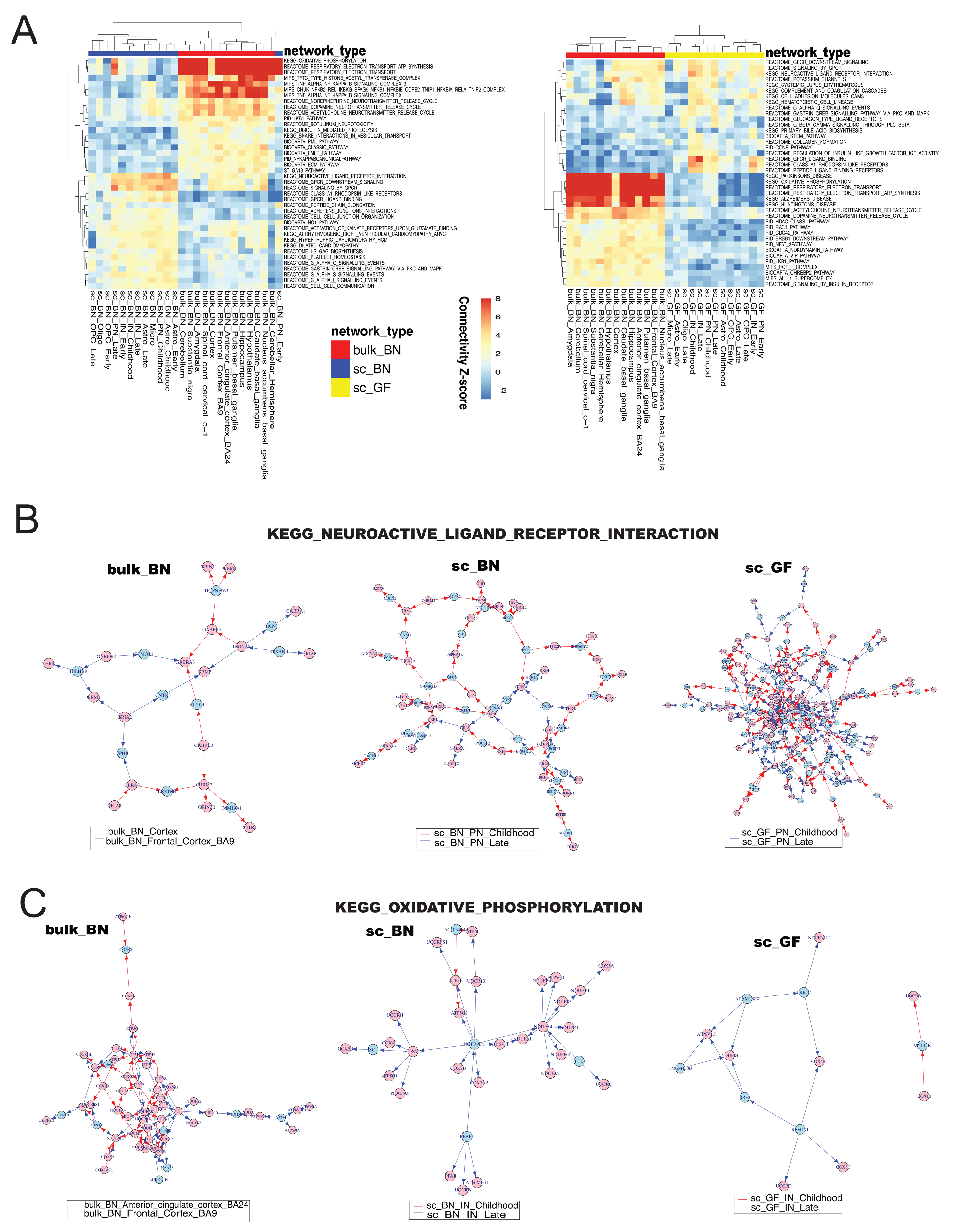
Differentially Connected Pathways between Bulk and Single-Cell Regulatory Networks. **(A)** Top differentially connected pathways between bulk_BN and sc_BN(left) or sc_GF(right). The connectivity Z-score (represented by color gradient) indicates the strength of connectivity within the pathways for each network. **(B)** Subnetworks of KEGG oxidative phosphorylation pathway (upper) and neuroactive ligand-receptor interaction pathway (lower) across the three network types: bulk_BN, sc_BN, and sc_GF. For each network type, two exemplary context-specific subnetworks were plotted together (distinguished by edge color). Red nodes represent genes in the respective pathway. Blue nodes are genes not within the pathways themselves, but highly connected to the red nodes (see Methods for details).

In summary, these network comparison results highlight the complementary nature of context-specific brain networks. Compared to bulk RNA-seq networks, scRNA-seq networks capture more variation in signaling pathways across different cellular states, such as among neuroactive ligand and receptor genes.

### Context-Specific Sub-networks Underlying Known ASD Risk Genes

We investigated the distribution of ASD risk genes across 39 context-specific brain networks. ASD risk genes were downloaded from SFARI database^12^, where genes were categorized into three groups based on confidence levels: high (232 genes), strong (702 genes), and suggestive (112 genes). First, we examined the relationship between ASD risk genes and hub nodes, finding that ASD risk genes were significantly enriched in hub nodes of multiple sc_BNs (Figure 4A). The most enriched network was sc_BN_IN_Early, suggesting that inhibitory interneurons, particularly those at early developmental stages (fetal, neonatal and infancy), are crucial contexts for investigating ASD gene mechanisms^13^. Notably, hub nodes of bulk_BNs of anterior cingulate cortex and ganglia tissues also showed enrichment for ASD risk genes, aligning with evidence that inflammatory changes affecting neuronal function in these brain regions may modulate social behaviors^14^.

**Figure 4:**
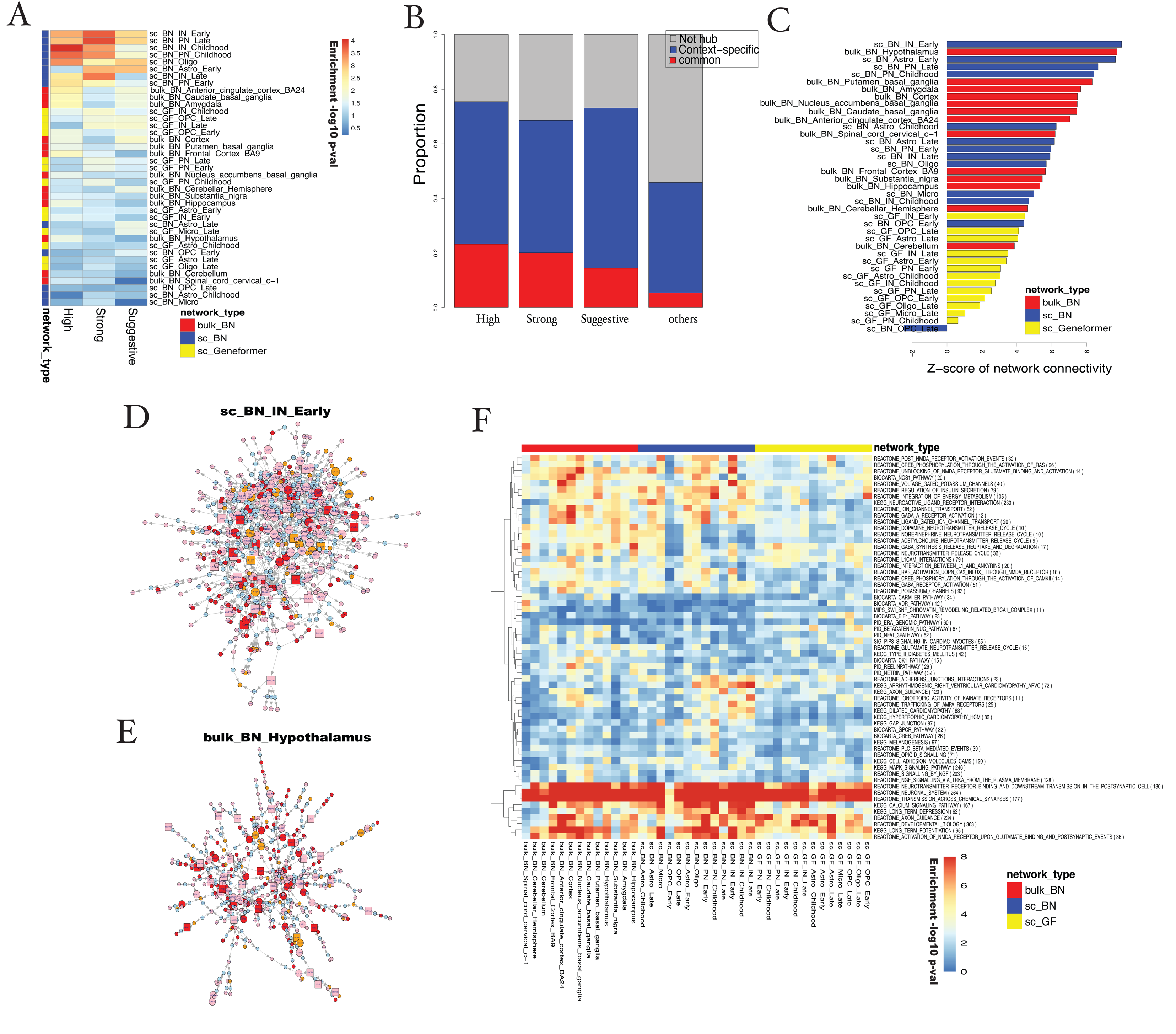
Distribution of Autism Spectrum Disorder (ASD) Risk Genes Across Bulk and Single-Cell Regulatory Networks. **(A)** Enrichment of high-confidence, strong and suggestive ASD risk genes in the hub nodes of each network. The color gradient represents the enrichment score calculated using a hypergeometric test (-log10 p-value). **(B)** The proportion of genes categorized as common hubs, context-specific hubs, or non-hub nodes, further stratified by their ASD risk level. **(C)** Network connectivity of ASD risk genes across the different types of networks. **(D)** Largest connected subnetwork for ASD risk genes in the sc_BN_IN_Early network (see Methods for details). Nodes in red, pink and orange represent high, strong and suggestive confidence ASD risk genes, respectively. Blue nodes indicate non-ASD risk genes. ASD risk genes that are hub nodes are larger than non-hub nodes, with round shapes indicating common hubs and square shapes representing context-specific hubs. **(E)** Largest connected ASD risk subnetwork in the bulk_BN_Hypothalamus network. The legend is the same as in panel (D). **(F)** Pathway enrichment results of ASD-risk subnetworks across different contexts.

Among common and context-specific hubs (as defined in above section), ASD genes were enriched in common hubs (Figure 4B). However, the majority of ASD genes were found in context-specific hubs, suggesting that these genes impact neuronal functions in context-specific ways, reinforcing the importance of constructing context-specific brain gene network to illuminate ASD mechanisms.

We next analyzed how known ASD risk genes are interconnected within these networks. Using the connectivity Z-score (as previously described), we found that ASD risk genes were significantly more interconnected compared to random gene sets of the same size in most networks (Figure 4C). Among the 39 networks, sc_BN_IN_Early had the highest connectivity, consistent with the hub node analysis, underscoring the critical roles of early-stage inhibitory interneurons in ASD. The largest connected subnetwork within sc_BN_IN_Early, underlying ASD risk genes (see Methods for details), includes 439 ASD risk genes (42% of all ASD risk genes) (Figure 4D). Gene set enrichment indicated that this subnetwork was enriched in genes related to synaptic transmission, including NMDA and AMPA receptors and calcium and potassium channels (a complete list is available in Table S2).

The second-ranked network by connectivity is bulk_Hypothelamus, with its ASD subnetwork shown in Figure 4E. Similar to the sc_BN_IN_Early subnetwork, the Hypothelamus subnetwork was also enriched in neuron transmission related pathways, albeit with weaker enrichment (Table S2). Certain gene sets were uniquely enriched in the Hypothelamus ASD subnetwork, such as the beta-catenin pathway, which is implicated in ASD at both genetic and transcriptional levels^15^. Our analysis suggests that the hypothalamus may be a critical context in which disruption of beta-catenin pathway causally contributes to ASD, consistent with this pathway’s essential role in hypothelamus development^16^.

We extended this analysis to all 39 networks and obtained 39 context-specific ASD subnetworks. Figure 4F (Table S2) displays the gene sets enriched in each subnetwork. Besides neuronal transmission and synaptic pathways, which were commonly enriched and are well-known to be related to ASD, some pathways showed context-specific enrichment. For example, cardiomyopathy-related pathways were enriched in IN/PN-specific subnetworks. A previous study highlighted an association between congenital heart disease (CHD) and ASD^17^, and ASD has also been linked to reduced heart rate variability^18^ and lower cardiac parasympathetic activity^19^. Such enrichment was exclusive to subnetworks derived from sc_BN, not bulk_BN or sc_GF, further emphasizing the importance of context-specific networks for exploring disease mechanisms.

### Context-Specific Key-regulators Underlying Differentially Expressed ASD Genes

In addition to genetic approaches, recent bulk and single-cell transcriptome analyses have provided complementary insights into the core molecular pathways underlying ASD. For example, the largest bulk RNA sequencing dataset of autism brains to date includes 47 autism and 57 control samples (ages 2-82) covering multiple cortical regions^20^. A recent study utilized single-nucleus RNA sequencing (snRNA-seq) to profile tissue samples from the prefrontal cortex and anterior cingulate cortex of 15 ASD patients and 16 controls (ages 4-22), generating cell type-specific differentially expressed genes (DEGs)^21^.

A major challenge in DEG analysis is distinguishing downstream genes from upstream causal key regulators, a task in which causal gene networks play an essential role^7,8^. We leveraged our 39 context-specific brain networks to identify key regulators for the DEGs from the two studies mentioned above. Specifically, we defined key regulators as hub nodes with network neighbors (within two steps) enriched for DEGs (adjusted p-value of fisher’s test <0.05). Figure 5A shows the number of key regulators identified for each DEG list(rows) across networks (columns). Interestingly, DEGs from bulk RNA-seq profiles yielded significantly more key regulators in bulk_BN networks than in sc_BN or sc_GF networks. Similarly, most key regulators for DEGs based on snRNA-seq profiles were identified in sc_BN and sc_GF networks rather than in bulk_BNs. The striking difference underscores the advantage of using context-matched gene networks to identify key regulators for DEGs.

**Figure 5:**
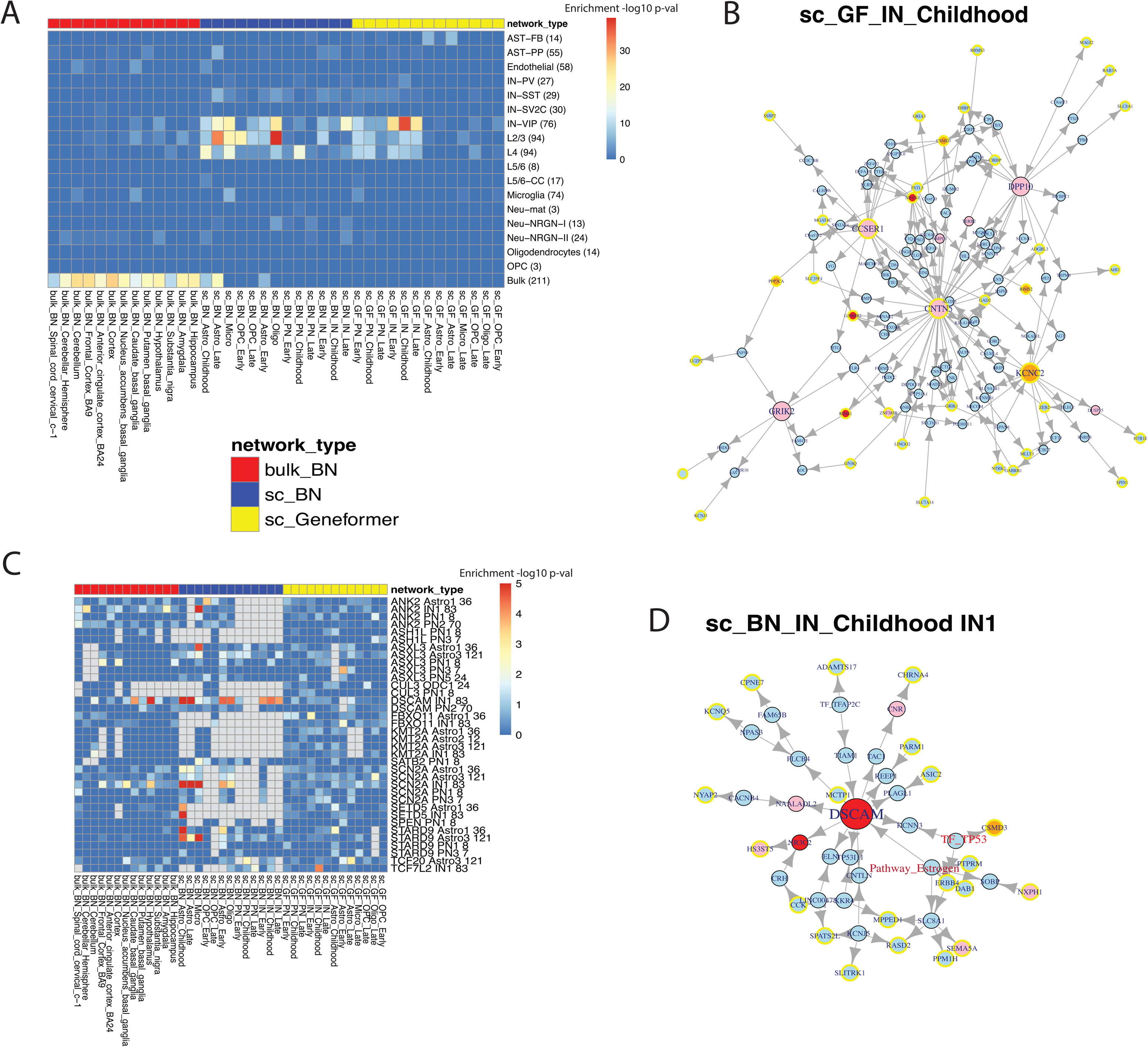
Integrating Context-Specific Networks with Differential Gene Expression in ASD and Perturb-Seq Results of ASD Risk Genes. **(A)** Heatmap showing the number of key-regulators identified across context-specific networks (columns) for differentially expressed genes (DEGs) in various cell types or bulk tissue of ASD patients (rows). The color gradient represents the count of key regulators identified for each combination. **(B)** Subnetwork of the top 5 key-regulators and differentially expressed genes detected in IN-VIP cells, extracted from the sc_GF_IN_Childhood network. Larger nodes indicate key-regulator genes. Nodes with yellow boundaries represent differentially expressed genes in IN-VIP cells. Red, pink, and orange nodes represent high, strong, and suggestive confidence ASD risk genes, respectively, while blue nodes represent non-ASD risk genes. **(C)** Overlap between network neighbors of ASD risk genes and differentially expressed gene modules observed in perturb-seq experiments. Each row represents an ASD risk gene and a cell type-specific gene module significantly perturbed by that risk gene in perturb-seq experiments. Columns correspond to context-specific networks. The color gradient shows the -log p-value of the hypergeometric test for the overlap between gene modules (rows) and the network neighbors of the ASD risk genes in the corresponding context-specific networks (columns). Grey color means the ASD gene is not included in that network. **(D)** Subnetwork of DSCAM and the perturbed IN1 module genes extracted from the sc_BN_IN_Childhood network. Nodes with yellow boundaries represent IN1 module genes. The legend of the node color is the same as panel (B).

The context-matching pattern can also be evident across cell types. For example, the network identifying the most key regulators for IN−VIP DEGs was sc_GF_IN_Childhood. For AST-FB DEGs, the top networks with the highest numbers of key regulators were sc_GF_Astro_Childhood and sc_GF_Astro_Late, while for AST-PP, the top network was sc_BN_Astro_Late. For Microglia DEGs, the top network was sc_BN_Micro. Note that these aforementioned top networks are at childhood or late stage instead of early stage, aligning with the ages of sample donors used to compute DEGs (ages 4-22). However, some mismatches were observed; for instance, the best networks for DEGS identified in different layers of PN were non-neuron networks from the sc_BN set. Overall, sc_GF networks appeared to better prioritize networks matching the cellular context of DEGs than sc_BN networks.

Given the critical role of inhibitory interneurons and the match between sc_GF_IN_Childhood network and the highest number of key regulators identified for IN-VIP DEGs, we further examined the subnetwork around the top 5 key regulators: "KCNC2", "CNTN5", "CCSER1", "DPP10", and "GRIK2", along with their neighbors (within two steps). (See Method for details, Figure 5B). The subnetwork included 36 out of 82 original DEGs. Notably, all five key regulators are known ASD risk genes, even though only 21% (17 of 82) of the original DEGs and 28% (10 of 36) of DEGs in the subnetwork are known ASD risk genes. This supports context-specific key-regulatory analysis as a potential approach for identifying crucial upstream disease-causing genes. Additionally, two ("DPP10" and "GRIK2") of the five key regulators were not DEGs themselves, indicating that key-regulator analysis can reveal disease genes that differential expression analyses might miss. The subnetwork was enriched in genes associated with chemical synapses and GABA signaling. For example, the top key-regulator “KCNC2” encodes a potassium channel essential for the function of GABAergic interneurons. Disruptions in GABAergic interneurons can lead to an imbalance between excitatory and inhibitory neurotransmission, contributing to ASD’s core symptoms^22^. Similarly, “GRIK2” encodes a member of the kainate receptor family, which are glutamate-gated ion channels that help maintain the brain’s excitatory and inhibitory balance^23^.

### Interpreting Perturb-Seq Results with Context-Specific Networks

To systematically study the function of ASD risk genes, a recent study^24^ used CRISPR-Cas9 to introduce frameshift mutations in 35 ASD risk genes within the developing mouse brain in utero, followed by scRNA-seq of major cell types to identify gene modules affected by each perturbation. Specifically, this study generated 14 cell-type-specific gene modules, associating each perturbed gene with gene modules in affected cell types^24^. Using an FDR cutoff of 0.1 for the association and requiring perturbed genes to be hub nodes in at least one of our 39 networks, we obtained 36 pairs of perturbed hub-node genes and their associated cell type-specific gene modules from the study. For comparison, we also obtained 39 pairs of perturbations and gene modules with FDR >0.7 as negative controls.

Our objective was to assess the capacity of our context-specific brain networks to recapitulate and interpret the 36 observed associations between gene perturbations and gene modules. We evaluated whether the affected gene module was enriched among the network neighbors of the perturbed gene (where network neighbors are defined as nodes within three steps). Figure 5C represents a total of 36*39 enrichment test results (-log p-value of Fisher’s exact test) for the 36 perturbation-gene module pairs (rows) across the 39 networks (columns). Grey indicates the perturbed gene was not included in the network. Despite the presence of missing genes, sc_BN networks best recapitulated context-specific perturbation responses (21 tests with nominal p-value <0.001 or FDR <0.07) compared to bulk_BN (4 tests with nominal p-value <0.001 or FDR <0.35) and sc_GF (3 tests with nominal p-value <0.001 or FDR <0.47). As expected, we observed much less enrichment when we performed similar evaluations on the 39 negative pairs of perturbations and gene modules (Figure S3, 0, 1, and 0 cases with nominal p-value <0.001 for bulk_BN, sc_BN and sc_GF networks, resepctively). There results support a significant concordance between perturb-seq results and our context-specific networks, especially with sc_BN networks. However, the concordance remains moderate, particularly when a matching cellular context is required. For instance, only 6 out of 21 enrichments detected in sc_BN networks were within matching cell types.

Among the six context-matching enrichments, three were related to DSCAM perturbation and an interneuron gene module (IN1), with sc_BN_IN_Childhood showing the strongest enrichment (p-value = 5.6e-5). We used this example to illustrate how our gene networks can enhance the interpretation of perturb-seq results. We extracted the subnetwork of DSCAM and IN1 module genes (See Method for details, Figure 5D). The DSCAM subnetwork was enriched with potassium channel and calcium signaling pathway genes (P-value=0.0017 and 0.0085), including CACNB4, CHRNA4, KCNJ5, KCNN3, and KCNQ5, with CHRNA4 and KCNQ5 being part of the IN1 module. This is consistent with the known roles of DSCAM in mediating synapse formation and neural development^25,26^.

Interestingly, the DSCAM subnetwork included a node representing the estrogen pathway activity. Numerous studies have suggested a protective role of estrogen in ASD^27-30^, and our network suggests that estrogen may exert its effects on ASD through DSCAM. Additionally, the subnetwork contained a node representing TP53 transcription factor activity. While TP53 is not listed among SFARI ASD risk genes, a recent study indicates p53 modulates activity-dependent synaptic strengthening and autism-like behavior^31^. Furthermore, another high-confidence ASD risk gene, NR3C2 (the Mineralocorticoid receptor), was directly regulated by DSCAM in the subnetwork. Although no literature currently supports a connection between these two high-confidence ASD risk genes, this finding suggests an avenue for future investigation.

## Discussion

This study highlights several critical differences between bulk and single-cell RNA-seq profiles, which in turn affect the brain gene regulatory networks we constructed. The bulk RNA-seq profiles consisted of over 100 subjects per tissue type, meaning that the bulk_BNs derived from these profiles reflect cross-subject covariations influenced not only by genetic regulation but also by confounding factors such as age, diet, environment, and cellular composition within tissues. In contrast, each set of cell-type and developmental stage-specific scRNA-seq profiles included no more than 10 subjects, allowing sc_BNs to capture co-regulatory dynamics specific to individual cell types within the same subject. Additionally, while the bulk RNA-seq profiles cover multiple brain regions, the scRNA-seq profiles are limited to the frontal cortex. Furthermore, bulk RNA-seq data was derived from older donors, while single-cell RNA-seq data came from much younger donors, suggesting that these networks may represent gene regulation at different developmental stages.

This specificity is critical for understanding cellular mechanisms in neurodevelopmental disorders such as ASD, as it allows for the identification of cell-type-specific regulatory hubs and signaling pathways that could drive disease processes. ASD is characterized by significant genetic heterogeneity and complexity. Deciphering the interplay between genetic, molecular, and cellular factors across brain regions and developmental stages is essential to uncovering the underlying causal mechanisms of ASD. Here, we constructed context-specific brain gene networks using both bulk RNA-seq and single-cell RNA-seq (scRNA-seq) data, applying integrative Bayesian Networks (BN) and the Geneformer deep learning model. Our analyses underscore the complementary value of these approaches in elucidating the multifaceted regulatory networks associated with ASD. Specifically, sc_BNs and sc_GFs showed better alignment with ASD differential expression and perturbation signatures derived from scRNA-seq data (Figure 5). Interestingly, sc_BNs and sc_GFs revealed higher connectivity among neuroactive ligand-receptor genes, which was less evident in bulk_BNs. This suggests that single-cell data more effectively captures context-specific signaling pathways, underscoring the importance of cell-type-specific regulatory mechanisms in ASD.

However, the lack of neurodegenerative disease gene enrichment in the scRNA-seq networks could be due to the younger age of donors in these datasets. Additionally, interactions between different cell types, detectable in bulk_BNs but not in sc_BNs or sc_GFs, may also contribute to the observed differences. These findings highlight the critical need for integration of bulk RNA-seq and scRNA-seq data to construct networks that capture inter-cellular communication and bridge insights from both bulk and single-cell networks.

Despite these findings, several limitations of this study must be acknowledged. Differences in sample size, tissue specificity, and age distribution between bulk and single-cell RNA-seq datasets could influence network construction and interpretation. Furthermore, while integrative BN has proven effective for gene regulatory network construction, further optimization is necessary for its application to scRNA-seq data, given the unique challenges posed by single-cell data, such as data sparsity and noise. Similarly, while deep learning models like Geneformer show promise for in-silico perturbation analysis, their interpretability and biological relevance require further validation.

The concordance between our networks and external datasets, such as perturb-seq results from scRNA-seq, was moderate. Specifically, we observed inconsistencies in cell type matches between affected gene modules and the most enriched networks. This discrepancy may result from limited biological context similarity between perturb-seq experiments (conducted in mice) and our network construction (based on human data). These interspecies differences underscore the complexity of translating findings from model organisms to human biology.

In conclusion, this study provides a comprehensive comparison of brain gene regulatory networks constructed from bulk and single-cell RNA-seq data, demonstrating their utility in understanding ASD. Our findings emphasize the complementary nature of multiple data modalities and computational approaches in capturing the context-specific gene regulations underlying ASD. This integrative framework enables the identification of key regulatory nodes and pathways that may inform targeted interventions for ASD. Future research should aim to systematically integrate bulk and single-cell RNA-seq data to reveal cell-type-specific gene regulations and their interactions with other cell types.

## Methods

### Construction of Integrative BN from Bulk RNA-seq

We downloaded genotypes, read count matrices, and pre-calculated eQTL analysis results for 13 tissues from various brain regions from GTEx release 8^1^ via dbGaP (accession number phs000424.v8.p2). Samples with paired RNA-seq and genotype data were selected for further analysis. The sample size per tissue ranged from 114 to 209, with a median of 170. RNA-seq profiles from tissues in similar brain regions showed clustering patterns (Figure S2A).

To construct Bayesian networks (BN) for each tissue dataset, we selected a consistent number of top variable genes. For each dataset, we computed the mean and standardized variance of each gene using the HVFinfo() function in the Seurat R package. Genes with the top 13,000 mean expression and top 8,000 standardized variance were initially selected. To increase the representation of disease-related genes, we reduced these cutoffs by 20% for genes known to have mutations associated with diseases^32,33^.

Knowledge-derived priors for sub-structures *P*(*S*|*K*) were assessed based on the likelihood equation *P*(*S*, *D*|*K*) = *P*(*S*|*K*)*P*(*D*|*S*, *K*), where *S* represents a sub-structure of interest, *D* is the data, and *K* is the knowledge collection. For deriving priors based on known human protein complexes, core protein complex data were downloaded from CORUM^34^. Tissue-specific co-expression patterns were used to evaluate the likelihood *P*(*D*|*S*, *K*). In brief, weak priors were set for pairwise gene-gene interactions unless a protein complex’s components showed significant transcriptional co-regulation (FDR < 1e-6 under random permutation). For each co-regulated complex, the member most highly correlated with other components was chosen as a surrogate regulator, setting a strong prior for a sub-structure with directed edges from the surrogate regulator to other members. Only large, co-regulated protein complexes with more than five directed edges received strong priors.

Similarly, priors were derived from databases of known transcription factor (TF)-target pairs and pathway regulations. TF and pathway activities were estimated using the Python package Decoupler^35^. TF activity was inferred based on a curated collection of TFs and their targets, including only those with confidence levels A and B. For each pathway, only the top 100 most significant targets were considered. Strong priors were set for sub-structures representing significant TF-target and pathway-target regulations, introducing TF activity as a regulator for target genes. This same approach was applied to integrate pathway activity with its targets.

Tissue-specific eQTL results from GTEx release 8 were further filtered to select significant cis-regulatory effects. Specifically, we required a nominal p-value of less than 1e-8, minor allele frequency (MAF) greater than 0.1, and log2 allelic fold change (aFC) greater than 0.5. Additionally, for each tissue, we included only significant eQTLs corresponding to the selected top variable genes. These eQTLs were incorporated as individual nodes in the network, with strong priors set for regulatory edges from the eQTLs to their cis-regulated target genes.

The gene expression and cis-eQTL nodes were then imported into the software suite, Reconstructing Integrative Molecular Bayesian Network (RIMBANet), to construct a biological causal network given the data and priors, as previously described^11,36^. Briefly, the network reconstruction process searches for a directed acyclic graph (DAG) structure *G* and associated parameters Θ that can best explain the given data *D*, *P*(*G*, Θ|*D*). If the structure *G* is a DAG, then *P*(*G*, Θ|*D*) can be decomposed into a series of sub-structures *P*(*G*, Θ|*D*) = ∏*_i_ P*(*G^i^*, Θ*^i^*|*D*). To speed up the searching process, for each gene, the bottom 20% genes based on their mutual information were excluded as potential candidate regulators (sparse candidate search^37^. The network reconstruction process is a Monte Carlo Markov chain (MCMC) process. We ran 1000 independent MCMC processes based on 1000 random seed numbers that resulted in 1000 candidate structures. Then, we selected consensus structure features with posterior probabilities >0.3 among candidate structures^38^. Finally, loops in the consensus network were removed by deleting the weakest link in the loops.

### Construction of Context-Specific Integrative BN from Single-Cell RNA-seq

Single-nuclear RNA-seq(snRNA-seq) data of the frontal cortex, spanning a broad age range from gestation to adulthood, were obtained from a previous study^3^. Specifically, the normalized and down-sampled data, consisting of 154,748 cells and 27 samples, were downloaded from http://brain.listerlab.org. Six neuron cell types (PN, IN, Astro, Oligo, OPC, and Micro) were included in the analyses. Cells were further subdivided based on donor developmental stage, balancing the stage-specific differences and sample sizes for each group. Specifically, PN, IN and Astro cells were divided into three developmental stages: Early (fetal, infancy and neonatal), Childhood and Late (adolescence and adult). The OPC cluster was split into two developmental stages (Early and Late), while Oligo and Micro were not further split due to limited sample size and minimal age-related variation. Overall, we grouped cells into 13 cell-type and developmental stage-specific clusters.

To address data sparsity issues inherent in snRNA-seq profiles, we merged single-cell profiles in similar cellular states into meta-profiles using R package MetaCell^39^. Parameters were adjusted to generate300-400 meta cells for each of the 13cell-type and developmental stage-specific clusters (details on the original single cells and derived meta cells are in Table S3). The approach ensured a similar number of meta-cell profiles across cell-type and developmental stages, minimizing the impact of sample size discrepancies on network construction accuracies. A umap plot (Figure S2B) illustrates clustering of these meta-cell profiles according to cell type and developmental stage.

Following the creation of meta-cell profiles, the remaining steps were similar to those in the construction of integrative BN from bulk RNA-seq profiles, including selecting top variable genes and integrating known protein complex, TF-regulation and pathway-regulation data in a context-specific manner.

### Construction of Context-Specific Causal Networks from Single-Cell RNA-seq using Deep Learning Model

In-silico perturbation was conducted using Geneformer^9^, a deep-learning model that was developed and pretrained on approximately 30 million single-cell gene-expression profiles, enabling it to make predictions about gene-network regulations. The codes and the pretrained model were downloaded from Hugging Face (repository: ctheodoris/Geneformer). To construct brain developmental stage-specific gene-networks, we fine-tune the pretrained model using the brain snRNA-seq data described above. Specifically, the model was fine-tuned with a cell classification learning objective, using the 19 major cell clusters annotated in the original snRNAseq dataset^3^.

For hyperparameter selection during fine-tuning, we split the scRNA-seq data into 80% for training and 20% for evaluation, using the HyperOptSearch() function from the Ray Tune Python package (<u>Ray Tune: Hyperparameter Tuning — Ray 2.30.0</u>) to optimize hyperparameters. The chosen hyperparameters included: a linear learning rate schedule with max learning rate of 5e-5, AdamW as the optimization function, a batch size of 12, weight decay of 0.001, and 10 epoch. As shown in Figure S2C, the model reached a plateau after just a few epochs. To illustrate the improvement in cell classification, we extracted cell embeddings from both the pretrained and refined models and plotted them in UMAP. Embeddings from the refined model were more consistent with the annotation of the 19 major cell clusters compared to those from the pretrained model (Figure S2D).

Using the refined model, we performed in-silico perturbation experiments with the InSilicoPerturber() function in Geneformer package. In a typical in-silico experiment, a gene (referred to as “source gene”) was “deleted” in a cell of interest, and the embeddings of all other genes (referred to as “target genes”) were computed before and after the perturbation. The cosine similarity of gene embeddings before and after perturbation was calculated for each target gene as a metric of the source gene’s effect on target genes. We repeated this experiment by perturbing each gene in each cell. Due to the time-intensive nature of this process, we down-sampled the snRNA-seq cell numbers to reduce computational costs. In this study, a maximum of 5 cells per major cell cluster per sample was selected to maintain cell heterogeneity, resulting in a total of 2,122 cells.

To construct the cell-type and developmental stage-specific gene networks from these in-silico perturbation experiments, we organize the cells into the same 13 cell-type and developmental stage-specific groups used in the BN construction from snRNA-seq. For each group, we summarized the effect of a source gene on a target gene by averaging cosine similarity scores across all cells in the group. We considered a source-target gene pair as a significant regulation if the averaged cosine similarity score was below 0.98 (approximately the top 2% of all possible gene pairs). The derived network includes all significant regulations, represented as directed edges from source genes to target genes. To avoid overly dense networks, each target gene was allowed a maximum of three source genes (those with the lowest cosine similarity scores). For edges in the final 13 networks, the cosine similarity score had a mean of 0.93 and a standard deviation of 0.049.

### Context-Specific sub-networks Underlying Known ASD Risk Genes

A total of 1,046 known ASD risk genes, encompassing all three confidence levels, were downloaded from SFARI database^12^. These genes were then mapped onto each of the 39 context-specific networks. Given that this list of ASD risk genes may not be exhaustive, we expanded it for each network by including nodes that were directly connected to at least two known ASD risk genes. The largest connected subnetwork associated with this extended set of ASD risk genes was extracted from each of the 39 networks and analyzed as the context-specific sub-networks underlying known ASD risk genes.

### Examples of DEG Key-Regulator Subnetwork and Perturb-seq Subnetwork

#### Key-regulator subnetwork around differentially expressed genes in IN-VIP cells

A list of differentially expressed genes(DEGs) between ASD patients and control samples, derived from the IN-VIP cell subpopulation, were downloaded from a previous study^21^ and mapped onto the sc_GF_IN_Childhood network. Key regulators were identified as hub nodes in sc_GF_IN_Childhood whose network neighbors (within two steps) were enriched with IN-VIP DEG list (adjusted p-value of Fisher’s test <0.05). For illustration, only the top five key regulators (ranked by p-value) were retained. Additionally, we identified mediator nodes between key regulators and DEGs (those within the shortest path between a key regulator and a DEG, with a maximum path length of 2). The largest connected subnetwork associated with DEGs, key regulators and mediator nodes was extracted from sc_GN_IN_Childhood network as an example of key-regulator subnetwork around IN-VIP DEG list.

#### Perturb-seq subnetwork between DSCAM perturbation and IN1 gene module

The IN1 gene module, downloaded from a previous study^24^, was mapped onto the sc_BN_IN_Childhood network. We identified mediator nodes between DSCAM and IN1 module genes (those within the shortest path between DSCAM and IN1 module genes, with a maximum path length of 3). The largest connected subnetwork associated with DSCAM, IN1 module genes, and mediator nodes was extracted from the sc_BN_IN_Childhood network as an example of a perturb-seq subnetwork illustrating the relationship between DSCAM perturbation and IN1 gene module.

## Supplementary Figure Legends

**Figure S1: Overview of Network Topology Across Bulk and Single-Cell RNA-seq Networks.**

**(A)** Distribution of node degree in each of the 39 context-specific networks. The x-axis represents node degree, and the y-axis shows the number of nodes with the respective degree.

**(B)** Boxplot of the average clustering coefficient of each network grouped by the network types.

**(C)** Frequency of hub nodes shared across networks.

**(D)** Heatmap clustering common hub genes into three groups, based on their presence in specific networks. Each row represents a hub gene, and each column corresponds to a context-specific network. Red indicates the gene is a hub node in that network, while blue indicates it is not. Rows are clustered using K-means clustering (K=3).

**(E)** Network similarity calculated by correlation coefficients for gene connectivity (measured by shortest path length) within each network.

**Figure S2: Overview of Data and Fine-Tuning of Geneformer model**

**(A)** UMAP plot showing the clustering of bulk RNA-seq data across 13 brain tissues, including cortex, cerebellum, and various subregions of the basal ganglia and spinal cord. The different tissues are color-coded.

**(B)** UMAP plot displaying the clustering of metaCell profiles derived from single-cell RNA-seq data, colored by six major cell types (left) or by developmental stages (right).

**(C)** The prediction accuracy for brain cell subpopulations improves and reaches a plateau after a few epochs of fine-tuning the original Geneformer model

**(D)** UMAP plot of scRNAseq profiles before and after parameter fine-tuning, with different cell subpopulations color-coded for visualization.

**Figure S3: Overlap Between Network Neighbors of ASD Risk Genes and Non-Significantly Perturbed Gene Modules in Perturb-Seq Experiments.**

This figure illustrates the degree of overlap between the network neighbors of ASD risk genes and gene modules that were not significantly affected in perturb-seq experiments. The legend is consistent with that of Figure 5C.

