## Supplementary figures and images for "Multiple Types of Context-Specific Brain Causal Regulatory Networks and their Applications to Autism Spectrum Disorder"

### Figure S1

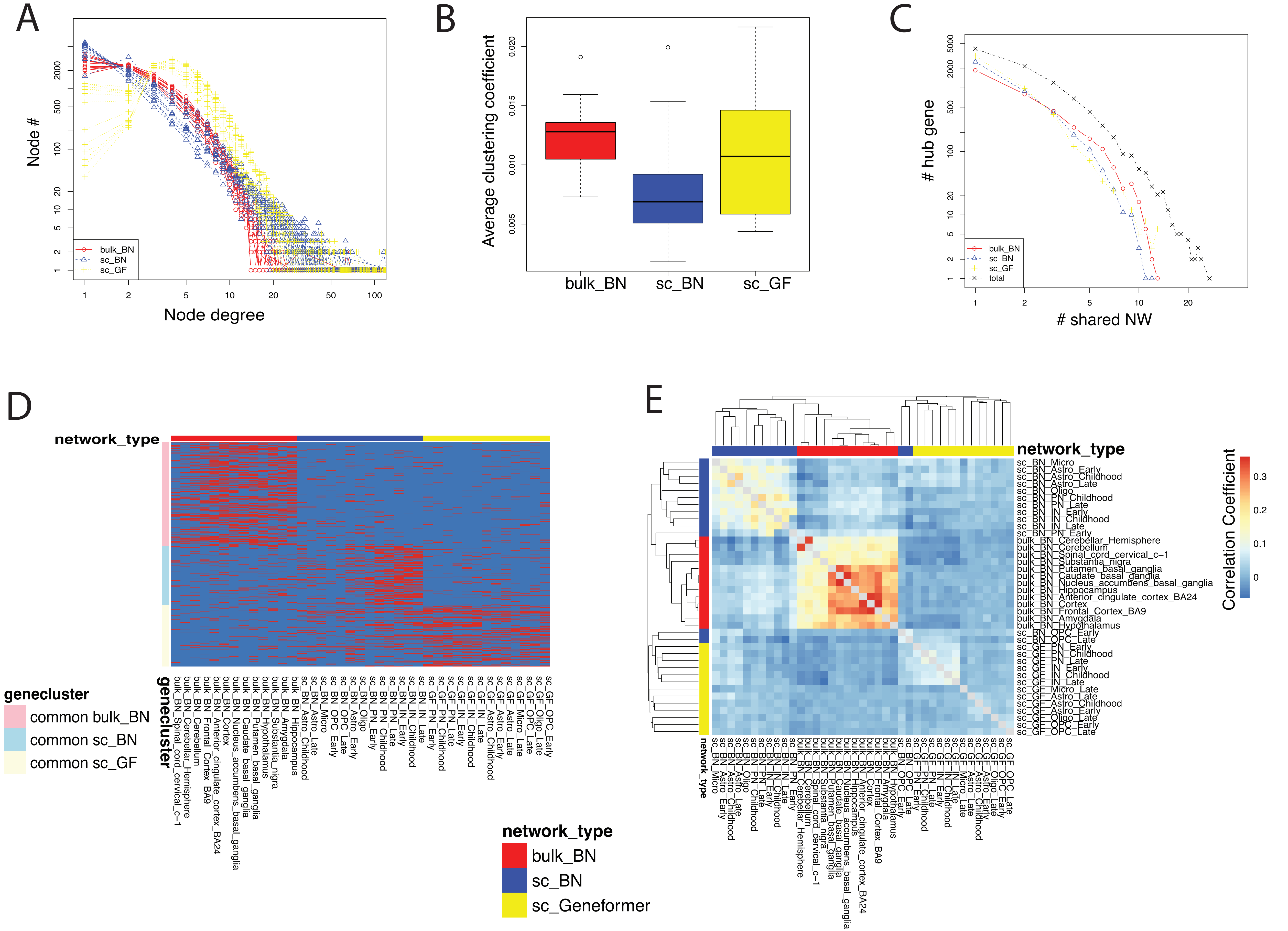

### Figure S2

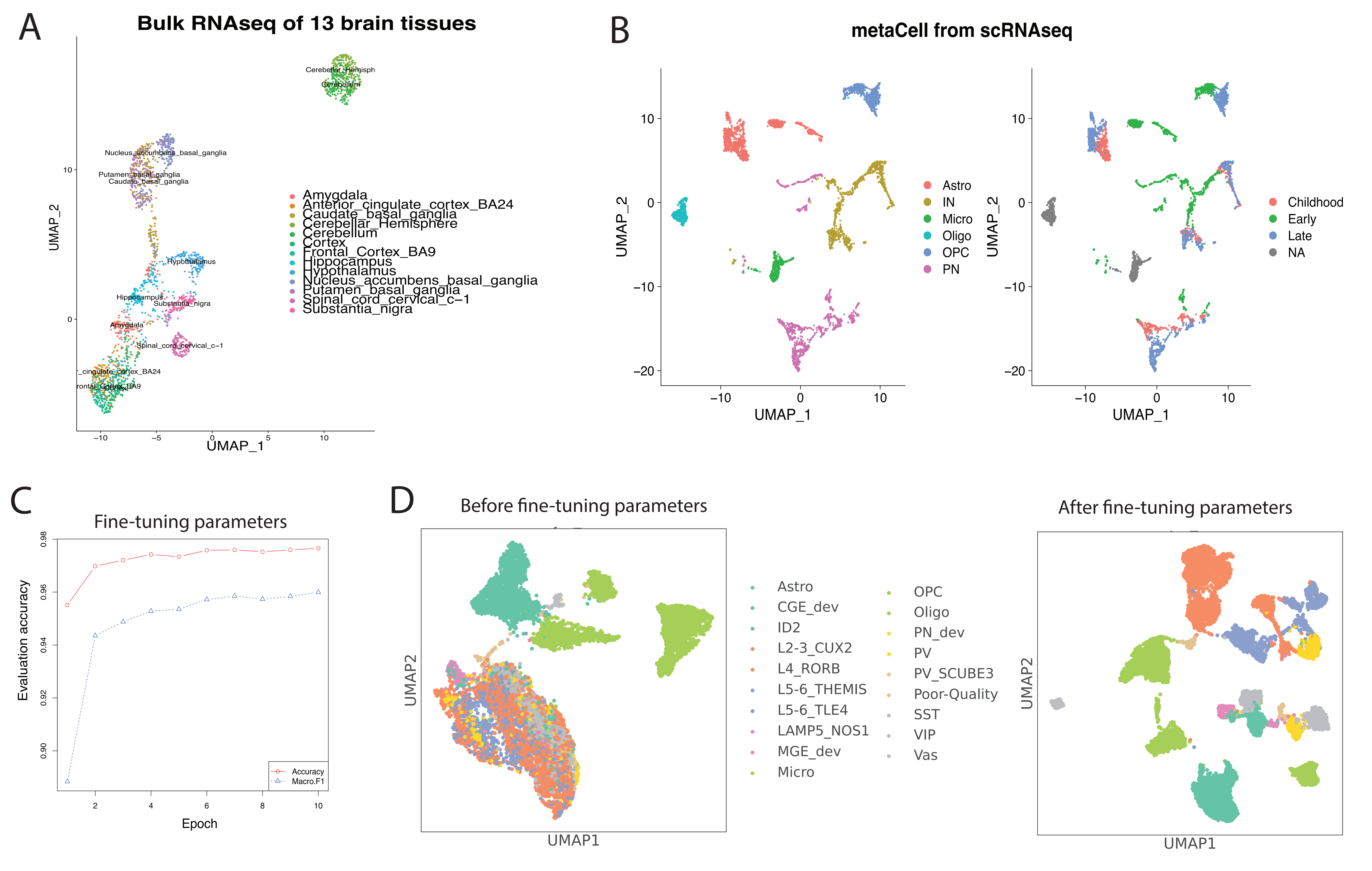

### Figure S3

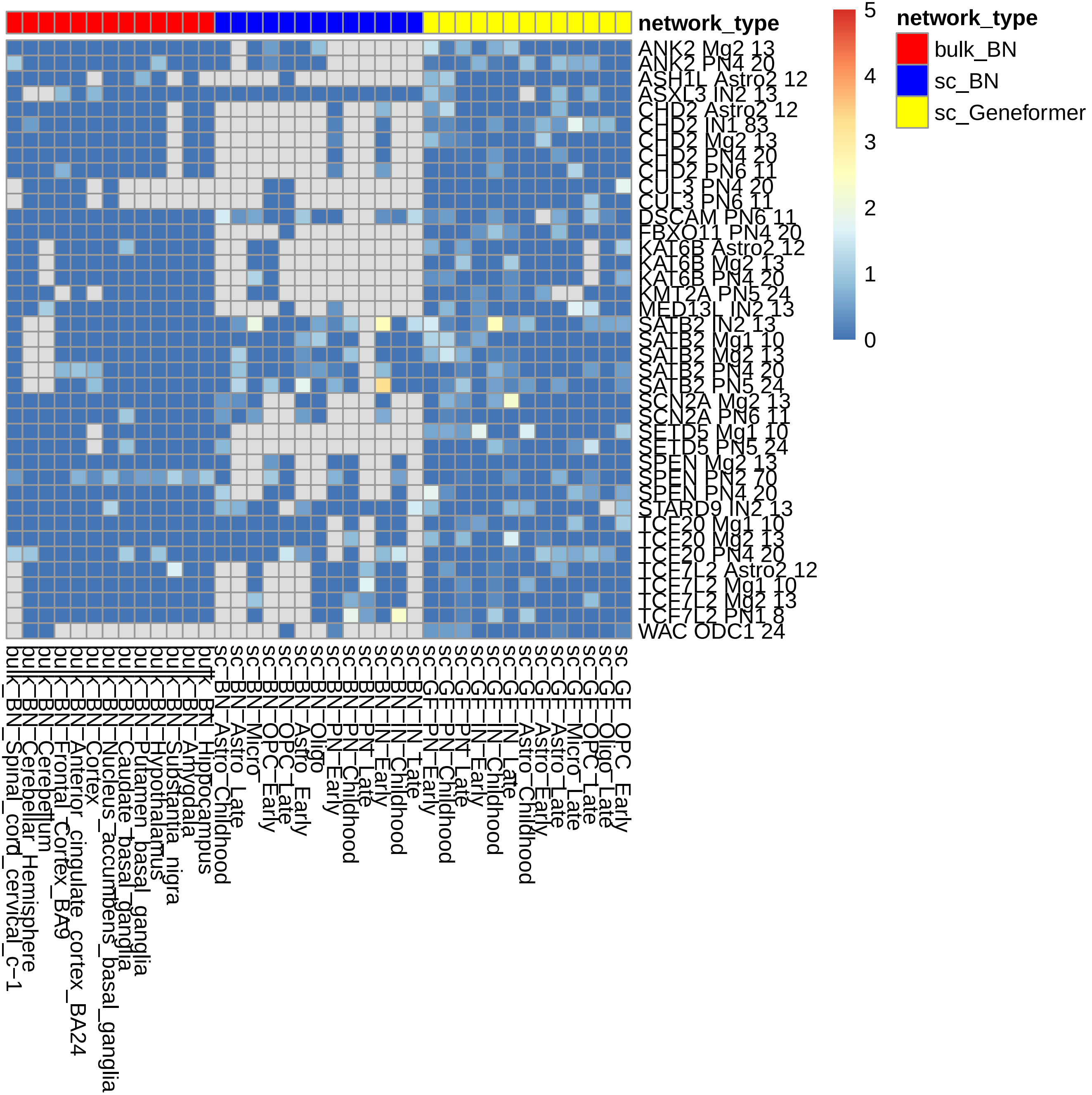
